# The influence of truncating the carboxy-terminal amino acid residues of Streptococcal enolase on its ability to interact with canine plasminogen

**DOI:** 10.1101/442004

**Authors:** Sasmit S. Deshmukh, M. Judith Kornblatt, Jack A. Kornblatt

## Abstract

The native octameric structure of streptococcal enolase from *Streptococcus pyogenes* increasingly dissociates as amino acid residues are removed one by one from the carboxy-terminus. These truncations gradually convert native octameric enolase into monomers and oligomers. In this work, we investigated how these truncations influence the interaction between Streptococcal enolase and canine plasminogen. We used dual polarization interferometry (DPI), localized surface plasmon resonance (LSPR), and sedimentation velocity analytical ultracentrifugation (AUC) to study the interaction. The DPI was our first technique, was performed on all the truncations and used one exclusive kind of chip. The LSRP was used to show that the DPI results were not dependent on the type of chip used. The AUC was required to show that our surface results were not the result of selecting a minority population in any given sample; the majority of the protein was responsible for the binding phenomenon we observed. By comparing results from these techniques we identified one detail that is essential for streptococcal enolase to bind plasminogen: In our hands the individual monomers bind plasminogen; dimers, trimers, tetramers may or may not bind, the fully intact, native, octamer does not bind plasminogen. We also evaluated the contribution to the equilibrium constant made by surface binding as well as in solution. On a surface, the association coefficient is about twice that in solution. The difference is probably not significant. Finally, the fully octameric form of the protein that does not contain a hexahis N-terminal peptide does not bind to a silicon oxynitride surface, does not bind to a Au-nanoparticle surface, does not bind to a surface coated with Ni-NTA nor does it bind to a surface coated with DPgn. The likelihood is great that the enolase species on the surface of *Streptococcus pyogenes* is an *x*-mer of the native octamer.

## Introduction

In 1991 Miles, Plow and their colleagues established that human plasminogen would bind to enolase on human cells [1]. Since that time, there has been a proliferation of the literature showing that enolase on a myriad of different cells will bind plasminogen. The enolase from *Streptococcus pyogenes* (Str. enolase) [2] (Fig 1A), as well as enolases from other bacterial sources, acts as a cell surface receptor for plasminogen [3,4]. The current view is that Str. enolase on the surface of the bacterial pathogen binds plasminogen, the plasminogen is activated to plasmin, and the plasmin helps to degrade some of the proteins which form the tight junctions between cells [5]. This promotes the spread of the pathogen [6].

Plasminogen is found in mammalian blood. The canine plasminogen, used in this work, consists of 793 amino acids organized into an N-terminal domain followed by five kringle domains and ending in a trypsin like domain (Fig 1B)[7,8,9]. Plasminogen is tightly knotted by 24 cystines.

**Fig 1:**
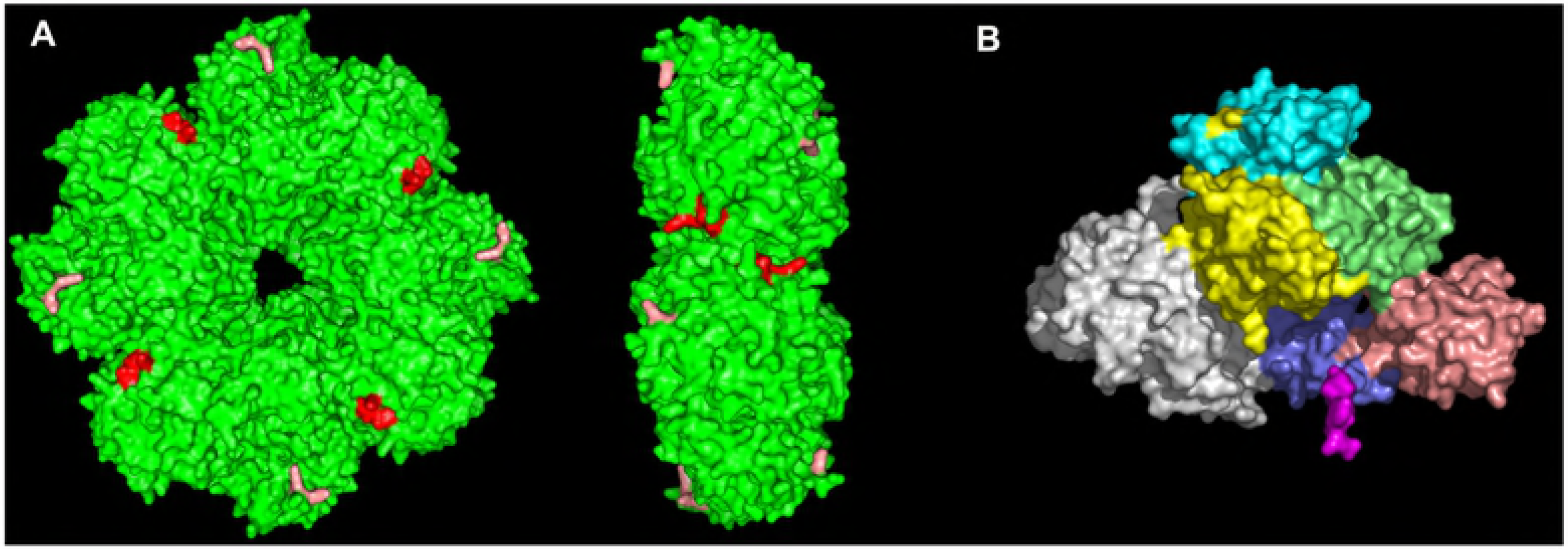
Surface representation of the Str. enolase and human plasminogen crystal structures. A. The homo-octamer of str. enolase (green) presented in front and side view. The red color represents the carboxy-terminal amino acid residues from 428 to 433 at the dimer-dimer interface where the truncations were made; the salmon color represents the amino acid residues 252 and 255. The last two residues, 434, 435, are not visible in the X-ray structure nor is the hexa-his tag. The dimensions of Str. enolase are 15 nm × 5 nm. B. Human plasminogen with the five kringle domains and the preproteolytic domain shown in different colors. Its dimensions are 10 nm × 8.5 nm × 5 nm. The crystal structures were taken from the PDB (Str. enolase 3ZLH) [10] and Plasminogen 4DUU [8]. These structures are not shown in identical scales.

Streptococcal enolase from *Str. pneumoniae* and the closely related Str. enolase from *Str. pyogenese* are homo-octamers[10–12]. Each monomer contains 435 amino acid residues. The octamer is tetramer of dimers (Fig 1A).

Each monomer of enolase has two proposed potential binding sites for plasminogen [11,12]. They are residues K435/K434 and residues K252/K255. 435 and 434 do not show in the X-ray structure. 433-428 are shown in red, while 252 and 255 are shown in orange in Fig 1A. Str. enolase was sequenced and the sequence deposited in the protein data bank (accession number NP_268959.1) [13]. The enolase clone used here was generously provided by Vijay Pancholi [14]; it has variations at F137L and E363G compared to the structure in the Data Bank (Str. enolase DB). We refer to the protein with its two changes as Str. enolase 137/363; it is the reference structure for all the work reported here.

The interaction between plasminogen and enolase in their native forms is not tight; it is sufficiently weak that we have not been able to detect it. However, when either protein goes from native to non-native, complex formation is easily detected [15,16].

The removal of carboxy-terminal amino acid residues close to the dimer-dimer interface of the enolase has been shown to destabilize the octameric structure of the protein and leads to monomers and oligomers [17]. Mutating residue 312 as well as other residues destabilizes the octameric structure and leads to reduced activity compared to the native octamer [10]. In this work we studied the interaction between plasminogen and mutated enolase species where the carboxy-terminal amino acid residues were removed sequentially from residue 435 to 428. We show that removal of the terminal amino acids does not stop Str. enolase from binding DPgn. The same is true when selected amino acids at the “second proposed binding site” are changed. When both proposed binding sites are removed, the enolase still binds plasminogen.

## Materials and Methods

The purifications and characterizations of canine plasminogen and Str. enolase have been described previously [16,18,19].

### Buffers and proteins

1. TME-S contains 50 mM Tris, 1 mM MgSO_4_, 0.1 mM Na_2_EDTA, 44 mN H_2_SO_4_ (TME-SO_4_) pH ~ 7.4). The octameric enolase is most stable in this buffer. All the AUC experiments were performed in this buffer. The enolase used in the DPI experiments was applied in this buffer.
2. Phosphate buffer 5mM KH_2_PO_4_ and 5 mM K_2_HPO_4_ at pH ~ 7 with or without 10 mM NaCl. This buffer was used for washing the silicon oxynitride chip in the DPI experiments. It destabilizes the octameric Str. enolase.
3. Pi, Imidazole, NaCl contained 5mM KH_2_PO_4_ and 5 mM K_2_HPO_4_, 10 mM imidazole, 10 mM NaCl. This buffer was used throughout the localizedSPR experiments.

Before use, all proteins were dialyzed for 48 hours with three changes of buffer. Following dialysis the proteins were centrifuged for 5 minutes (13 000 rpm, RT) to remove the largest aggregates. The influence of the three buffers on the structure of Str. enolase is significant, is dependent on the initial equilibrium position of the octamer/monomer equilibrium and is shown in Figs S1 and S2.

All chemicals were of highest purity available. Most were purchased from Sigma-Aldrich or Fluka Chemical Co.

The reference protein for the work reported here is Str. enolase F137L/E363G. It has an N-terminal hexa-histidine tag. We have also worked with the Str. enolase whose amino acid sequence is identical with that in the data bank (Str. enolase DB). It and all the mutated forms of Str. enolase have a hexa-his tag at the N-terminus **except Str. enolase DB no hexa-his tag**. Considering Str. enolase 137/363, removal of first carboxy-terminal amino acid residue at position 435 is referred as Str. enolase137/363 −1. Removal of the two carboxy-terminal residues is referred as Str. enolase 137/363 −2 and so on. For this study we used variants of Str. enolase 137/363 from which up to eight carboxy-terminal residues were removed. Site directed mutagenesis, expression and purification have been described elsewhere [17,19].

### Dual Polarization Interferometry (DPI)

Real time surface layer thickness and mass were measured using an *Ana*Light Bio200 dual polarization interferometer (FarField Group Ltd. Manchester, UK). The technique was described previously. The descriptions include those for studying protein-protein interactions, formation of thin films on the surface and structural changes at sub-nanometer scale[15,20–25]. This instrument is no longer supported by the company. Unmodified silicon oxynitride *Ana*Chip FB 80 chips (FarField Group Ltd. Manchester, UK) were used to measure the dimensions and protein-protein interactions in the adsorbed layers. The chip surface was extensively cleaned with 2% Hellmanex (Sigma-Aldrich) followed by isopropanol prior to each experiment. All measurements were done at 20 °C with a pump flow rate of 25 μL/min/channel. The plasminogen and Str. enolase protein concentrations were 3 μM each. Each protein was injected three times with injection times of 8 min so that the chip surface was close to saturated. The proteins were in TME-SO_4_. The wash buffer was 10 mM phosphate. Phase changes in terms of transverse magnetic (TM) and transverse electric (TE) values were fitted using *Ana*Light Explorer software to calculate the average thickness and mass of the adsorbed layers [15,26].

Both the original DPI data as well as the transformed data are included in Supplemental Files: all files for figures and tables\all dpi files. The files carry either an xpt or an xls suffix. The xls files are transformed from the xpt. The xpt files can only be read with the protected software. The xls files can be read with any program that accepts Excel files.

### Localized Surface Plasmon Resonance (LSPR)

We used a Nicoya Life Sciences OpenSPR instrument which operates in the localized SPR mode. The technique is not new; it has been successfully used to measure protein-protein interactions as well as protein-small molecule interactions. The principle of the measurements, as well as some examples, has been well described [27–33]. All experiments were performed with proteins that had been dialyzed against Pi, imidazole, NaCl. The running buffer for the experiments was the same as that for the dialyses. The experiments included those on Au-chips as well as those on Ni-NTA derivatized chips. The LSPR was not thermostated but the temperature of each run was close to 27 °C.

### Sedimentation Velocity Analytical Ultracentrifugation (AUC)

Solution phase binding of the Str. enolase and plasminogen was performed on a Beckman XL-I instrument running at 20 °C at 37000 rpm [34]. The resulting sedimentation data were evaluated using Sedfit c(s) version 14.1 [35,36]; Sedphat ver 12.1 was used to evaluate a K_A_ for the 137/363 -4/DPgn interaction [37–39].

Plasminogen alone, Str. enolase alone and the mixture of two proteins at the same concentrations as in the single protein samples were included in each run to determine if the s value of the plasminogen shifted in the presence of Str. enolase. The functional concentration of the two proteins in the mixture cell was established by integrating the individual peaks in the single protein samples, and then multiplying the % of the absorbance by the added concentration of total protein. For DPgn which is close to 95% pure this resulted in a very small correction. For 137/363 -4, which is between 80% and 90% pure but is polydisperse, the calculated functional concentration for the monomer peak was considerably less than the added total protein.

We used Sedfit to qualitatively establish whether there was an interaction between the two proteins. c(s) easily handles large data sets; it is extremely precise and is model dependent. Differences in s values greater than 0.5% are real.

We have also used DCDT+ (DCDT+ version 2.4.3 [40–42]) to evaluate the AUC data. The DCDT+ approach is discussed in the supplementary text 1.

As previously reported, the truncation of Str. enolase 137/363 enolase from the carboxy-terminal resulted in a gradual dissociation of the homo-octamer into monomers and oligomers [17].

The original AUC data are included in “all files for figures and tables” and the sub-files found therein.

## Results and Discussion

We were motivated to undertake this work by our finding that truncation of the carboxy-terminus of Str. enolase 137/363 led to more and more dissociation as the residues were removed one by one. We can easily detect DPgn binding to Str. enolase 137/363 on solid surfaces or on aggregates of either species. We have never tested Str. enolase 137/363 binding in solution to DPgn when the former was dissociated. The C-terminal amino acids of Str. enolase are located near the dimer-dimer interface. Our earlier work showed that removal of up to eight of these amino acids destabilized the octameric structure and shifted the equilibrium

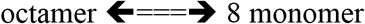
in the direction of monomers [17]but also gave rise to multimeric species. The shift resulted from a gradual destabilization of the interface as well as potential rearrangement of the packing of the monomers. This destabilization was quantified and was shown to be only ~half a kcal per monomer per residue removed. This was enough to cause noticeable shifts in the above equilibrium which could be easily determined by loss of activity and by analytical ultracentrifugation.

As stated, each monomer of Str. enolase contains two well characterized potential binding sites for plasminogen. The first is associated with the two C-terminal lysines: residues 435 and 434 [2,43]. The second consists of two internal residues: lysines 252 and 255[14,11,44]. The area close to residues 434 and 435 is highlighted in Fig 1A in red, the internal residues are in salmon. The actual C-terminal residues do not appear in the X-ray structure. By analogy with antibody-antigen interactions, the total binding site on the Str. enolase must be more extensive than two lysines. For instance, when comparing the cross reactivity of a monoclonal antibody raised against a synthetic peptide of sperm whale myoglobin, it was found that other myoglobins, even those with the identical sequence, had different reactivities with the monoclonal antibody[45]. This could have been due to different percentages of heme inserted correctly into the heme pocket but this was not discussed by the authors. Would we be able to detect binding of these truncated and mutated species of Str. enolase to DPgn in either solution or the solid phase? We address this question in the remainder of this work.

Can we detect binding of DPgn to the immobilized, truncated and mutated variants of Str. enolase on a solid surface? The answer is an unequivocal yes (Fig 2, 3).

The binding data in Figures 2, 3 and Table 1 are based on dual polarization interferometry measurements. DPI sensorgrams are analogous to those of surface plasmon resonance (SPR). DPI uses two polarized light beams to determine refractive indices which allows for the calculation of layer thickness on the chip. DPI allows us to characterize protein-protein interactions and structural changes that might arise from those interactions [21]. Since this is a surface technique, the orientation of the molecule under study is important for specific binding studies: specific binding sites must be exposed. Where binding is either non-specific or a mix of specific and non-specific, orientation becomes less stringent. Fig 2 is shown for illustrative purposes only; it shows the nature of the raw data sensorgram. The ordinate is radians. The scales in Fig 2A and 2B differ by a factor of 60 but the dynamic range of the instrument is considerably greater than shown in the figure. Fig 2A shows that not all proteins bind to the chip. Str. enolase DB with no hexa-his tag does not bind (2A, Table1). Fig 2B shows a sensorgram in which binding to the chip is rapid; the first three injections are Str. enolase 137/363. It binds and saturates the chip; the Str. enolase 137/363 is followed by two injections of DPgn which saturates the binding sites on the enolase. Fig 2A allows us to draw an important conclusion: *A portion of the binding energy of Str. enolase to the silicon oxynitride chip is contributed by the hexa-his tag. This 20 residue addition to a protein is not necessarily benign*.

**Fig 2:**
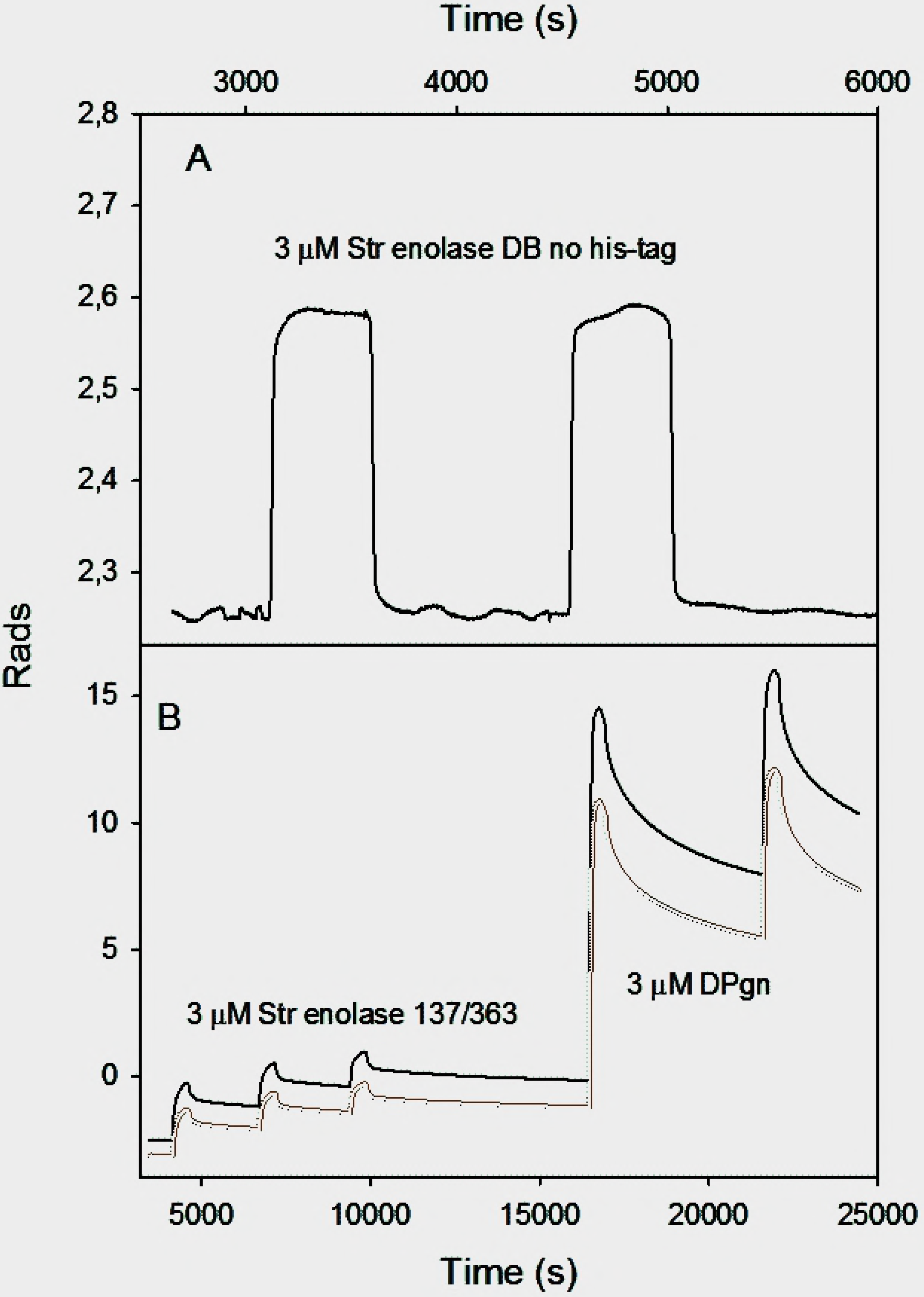
DPI, the raw data. **A**. The raw data of a DPI measurement representing deposition of Str. enolase DB with no hexa-his tag on a silicon oxynitride chip. Only one channel is presented. As shown in the last line of Table 1, this protein does not bind to the chip. **B**. Raw data from a single channel. The figure shows the detector response when three injections of Str. enolase 137/363 were made. This saturated the chip. The Str. enolase 137/363 was followed by two injections of DPgn which bound to the Str. enolase surface layer. The black, solid trace is the TM signal; the dotted trace the TE signal.

Based on data such as that in Fig 2B, Fig 3 shows the binding patterns for the truncated enolases as determined by DPI. The figure shows that all the enolase forms (except Str. enolase DB no his tag (Fig 2A)) bind DPgn when enolase is the first protein to bind to the chip. Inspection of the raw data indicated that there was neither enhanced binding nor decreased binding as we went from the ***majority*** of the protein being octamer to the majority of the soluble protein being monomer. The red circles show that when enolase is the first protein injected it binds to the chip. Neither the layer height nor the mass is dependent on the degree of truncation. This indicates that the binding (site) of the Str. enolase to the chip is not significantly perturbed by dissociation of the octamers to monomers and other species. When DPgn was subsequently injected, it bound to the Str. enolase layer. The total mass of protein that was bound was independent of the degree of truncation (Fig 3B). The total thickness of the Str. enolase plus DPgn appears to be dependent on the degree of truncation (Fig 3A). It has to be emphasized that in our hands, there is a significant degree of variation in both mass and thickness determinations. The variation of thickness as a function of degree of truncation shown in Fig 3A may be real but may also be the result of too few determinations. In a previous study [15] the total thickness of the Str. enolase 137/363 followed by DPgn layer was comparable to that shown here. We conclude from Fig 3B that all the Str. enolase 137/363 forms, −0 to −8, can bind DPgn.

**Fig 3:**
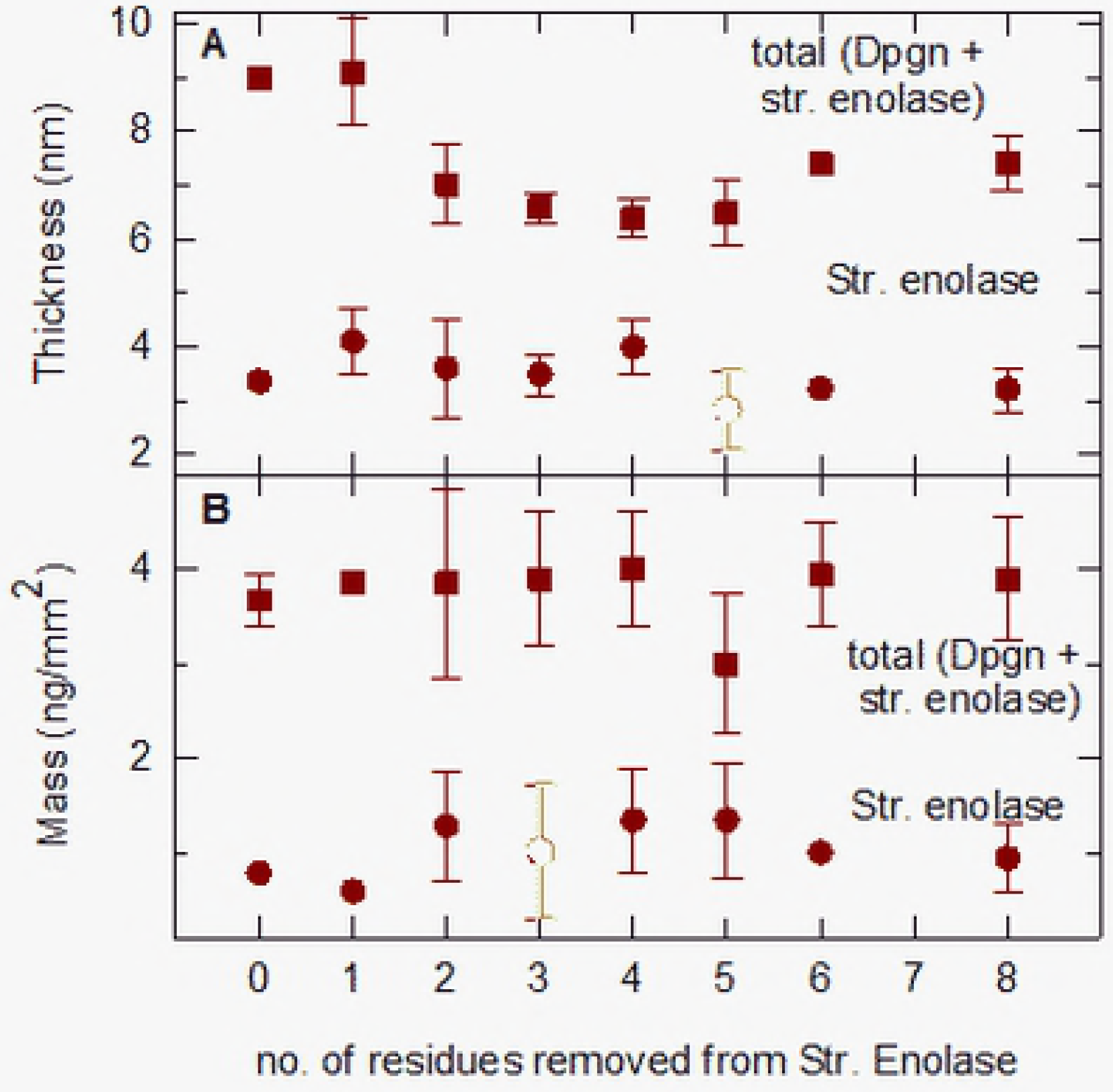
Analyzed data of DPI measurements. Thickness (Panel A) and mass (Panel B) resulting from the deposition of Str. enolase variants on the sensor chip are shown in the figure. These are represented by filled circles. The total thickness and total mass, as a result of subsequent deposition of DPgn on the pre-deposited Str. enolase variants, are represented by filled squares. The data shown are average of at least two experiments and the error bars represent variations in determining precise values from multiple experiments. For data where the variation is within the symbol, error bars are not shown.

The data of Table 1 show that on a surface, Str. enolase DB (with the his-tag) and Str. enolase 136/363-2 K252A/K255A still bind DPgn. Str.enolase DB, no his-tag does not bind to the chip; the DPgn that deposits binds to the empty chip. The Str. enolase 136/363-2 K252A/K255A result is particularly noteworthy since both well studied binding sites have been removed.

**Table 1.** DPgn binding to other forms of immobilized *Streptococcus pyogenese* enolase determined by DPI.

| Enolase form | % octamer<br>(from AUC) | % monomer<br>(from AUC) | Mass of<br>enolase layer<br>(ng/mm<sup>2</sup>) | Mass of<br>enolase +<br>DPgn layer<br>(ng/mm<sup>2</sup>) |
| --- | --- | --- | --- | --- |
| Str. enolase<br>DB | 88 | 0.4 (nil?) | 1.00 | 2.8 |
| Str. enolase<br>137/363 -2<br>K252A<br>K255A*** | Nil | 70.4 | 0.30 | 5.3 |
| Str. enolase<br>DB, no<br>hexahis tag | 81.6 | 1.3 | ~0.04 | 2.1 |

\*\*\* Str. enolase 137/363 -2 K252A K255A has had both proposed DPgn binding sites removed.

Dual polarization interferometry, and LSPR discussed in the next section, suffer from the same problems that all microfluidic techniques have. There is a limited area on the chip surface onto which proteins can adsorb or bind to proteins that are already present. The total amount of protein that is flowed over the chip is in vast excess compared to the total amount of protein that can bind. The ratio of applied protein to bound is > 1000. The result of this is that the protein that binds is selected. If the samples are 100.0% pure this is not an issue. We estimate that our proteins are between 80% and 95% pure which means that the chip surface can select out that 5%-20% of protein that binds more tightly than the bulk. The Str. enolase 137/363 is primarily octameric. The Str. enolase DB is for all intents and purposes octameric. The Str. enolase 137/363 truncations are a mixture of octamers and other species. The plasminogen is highly pure but not 100% pure. We do not select our data. However, the silicon oxynitride chip may select proteins that are not those of interest and thereby bias the analysis. A friend reading an early version of the manuscript indicated that it would be significantly stronger if it was supplemented by data taken with chips having other surface properties. Accordingly, we turned to surface plasmon resonance where such chips were available. We used LSPR with either Au nanoparticles or Ni-NTA surfaces as binding substrates.

The DB no tag will not bind to a gold nanoparticle chip (Fig S3) just as it does not bind to a silicon oxynitride chip; it will also not bind to an Au chip saturated with DPgn (Fig S4). In contrast to the DB no tag, Str. enolase 137/363 -4 does bind DPgn. Fig 4 shows the results obtained when Str. enolase 137/363 -4 is allowed to bind to a Ni-NTA chip and then titrated with DPgn. Str. enolase 137/363 -4 was applied continuously to the Ni-NTA chip at a rate of 20 μL per minute over a period of about 2 hours. The chip was then washed with buffer for another 2.5 hours before the application of 11 injections of DPgn. The injection schedule is shown in Supplementary text 2.

**Fig 4.**
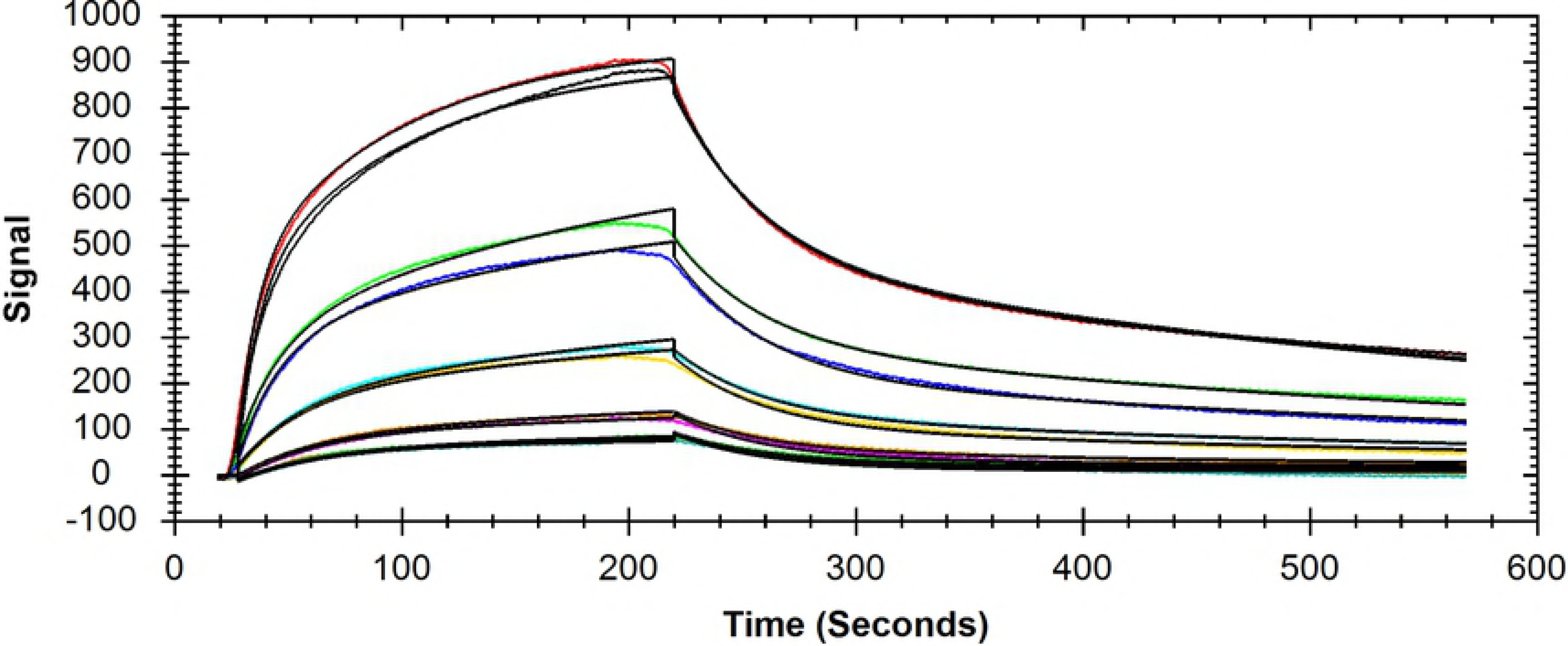
Binding of DPgn to Str. enolase 137/363 -4 immobilized on a Ni-NTA chip. Str. enolase 137/363 -4 was applied continuously to the Ni-NTA chip from second ~2000 to second 9000. At second 18000 DPgn injections were started. See supplementary text 2 for the injection schedule.

The data were evaluated for a dissociation constant using the program Trace Drawer (Version 1.5, Uppsala, Sweden). The fit is shown in Fig 5.

**Fig 5.**
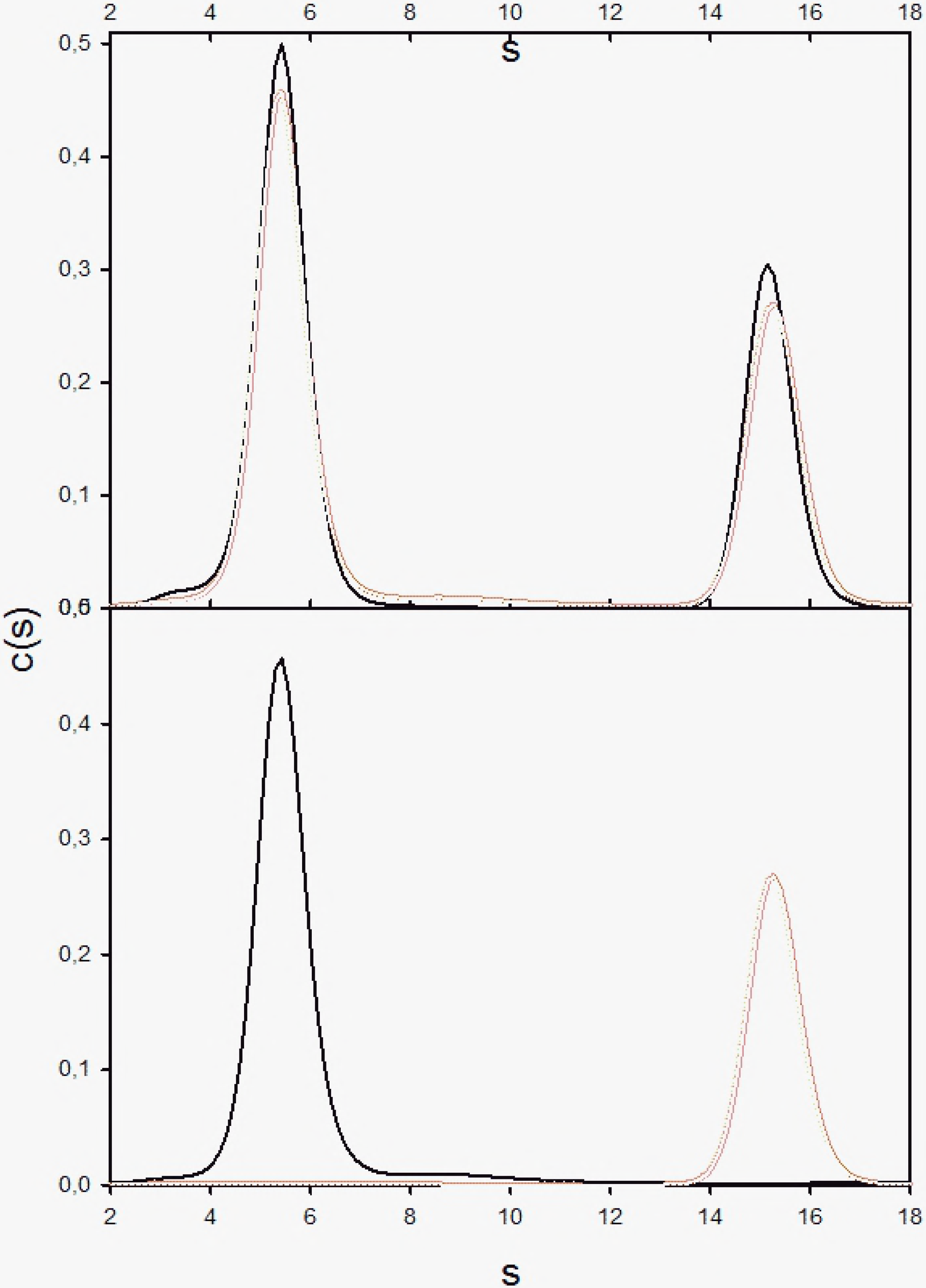
Analysis of the binding of DPgn to Str. enolase 137/363 -4 immobilized on Ni-NTA. A one to two model was fit to the data of Fig 4. The different injections of DPgn are shown as colored traces whereas the black lines are the fits. Each concentration was done in duplicate.

A one to two model was fit to the data. A one to two binding model means that the protein on the chip has two independent binding sites for the captured protein (Trace Drawer manual). The fitted thermodynamic parameters evaluated at a K_D1_ of 1.34 E-6 and K_D2_ of 2.66 E-6. Although the K_D_s are very close, the on and off rates for the two sites differ by almost an order of magnitude. The fits are very good but it must be emphasized that fits using a one to two model are always better than a one to one. In this instance K435/K434 has been removed; we know that there is at least one documented independent binding site on this Str. enolase (K252/K255) and the data imply a second. The chip, saturated with Str. enolase 137/363 -4 contained 1.5 E-12 mols/cm^2^. When then saturated with DPgn it contained 2.4 E-12 mols/cm^2^ of DPgn (Nicoya Lifesciences). The one enolase monomer to 1.6 DPgn is an approximate value but also justifies a tentative two binding sites in addition to the N-terminal K435/K434.

The above results from LSPR indicate that, overall, our DPI study yielded conclusions that were reliable. What were those conclusions? First, the intact Str. enolase octamer, without any tags, is not the species that binds plasminogen. Second, all full length species of Str. enolase that dissociate on time scales comparable to experiments are capable of binding plasminogen. Third, neither of the two proposed binding sites, the carboxy terminal dilysine and the internal site K252 and K255, are required for binding. We earlier emphasized that DPI and LSPR may look at minor components in a mixture.

In contrast to those techniques, AUC looks at all the protein in the sample except for that which has gone to the bottom of the cell prior to the centrifuge reaching speed. If there is a very minor component, a contaminant or a modified form of the major species, it may be selected for binding but it might not be noticed in the fitting routine. It follows then that, unless the proteins are 100.0% pure, there are significant differences between the conclusions that can be drawn from the two techniques. We have not in the past been able to detect interaction between the two when free in solution [15,16]. Would we be able to detect binding of the dissociated Str. enolase to DPgn?

The Str. enolase 137/363, the truncations, the Str. enolase 137/363 -2 K252A K255A and the Str. enolase DB were all subjected to sedimentation velocity analytical ultracentrifugation in the presence and absence of DPgn. In each case, the rotor contained one cell with pure DPgn, one with the pure Str. enolase and one cell with a mix of the two proteins at the same concentrations found in the pure protein containing cells. The protein distributions were evaluated using Sedfit c(s). We have also used DCDT+ to evaluate the AUC data.

The distributions of two sets of AUC data are shown in Figs 6 and 7.

In all the figures, DPgn alone runs at an s_20, w_ of close to 5.6. Our previous study indicated that the Str. enolase monomer runs at an s_20, w_ value of 3.2 [46] and the yeast enolase dimer runs at an s_20, w_ of 5.6 [47] The AUC data can be summarized rather easily. Whether or not we saw binding of a particular Str. enolase form to DPgn in the AUC was dependent on the amount of non-octamer present in the preparation. Str. enolase DB no tag, Str. enolase DB and Str. enolase 137/363 always contained a variable amount of non-octamer. We never saw binding of the DB no tag or the DB forms. We would occasionally see binding to the 137/363 form. The DB data are shown in Figs 6 and S5.

**Fig 6.**
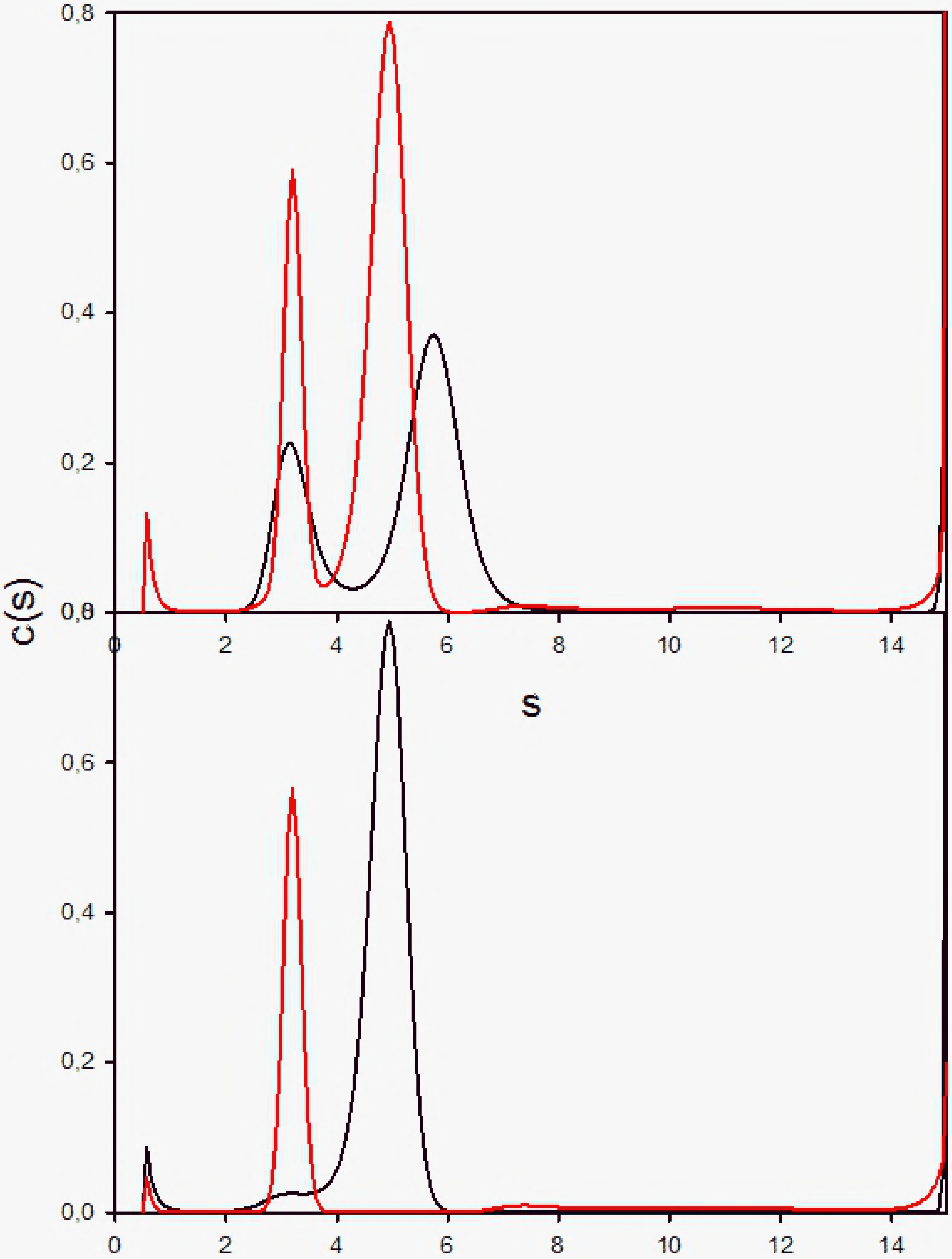
AUC of Str. enolase DB, DPgn and the mix of the two proteins. The Sedfit analysis. The top panel shows that the two proteins do not appear to interact in the analytical ultracentrifuge. The black trace is the c(s) fit to the actual mixture while the red trace is the sum of the fits to the individual proteins shown in the lower panel (black, DPgn; red, Str. enolase DB).

**Fig 7.**
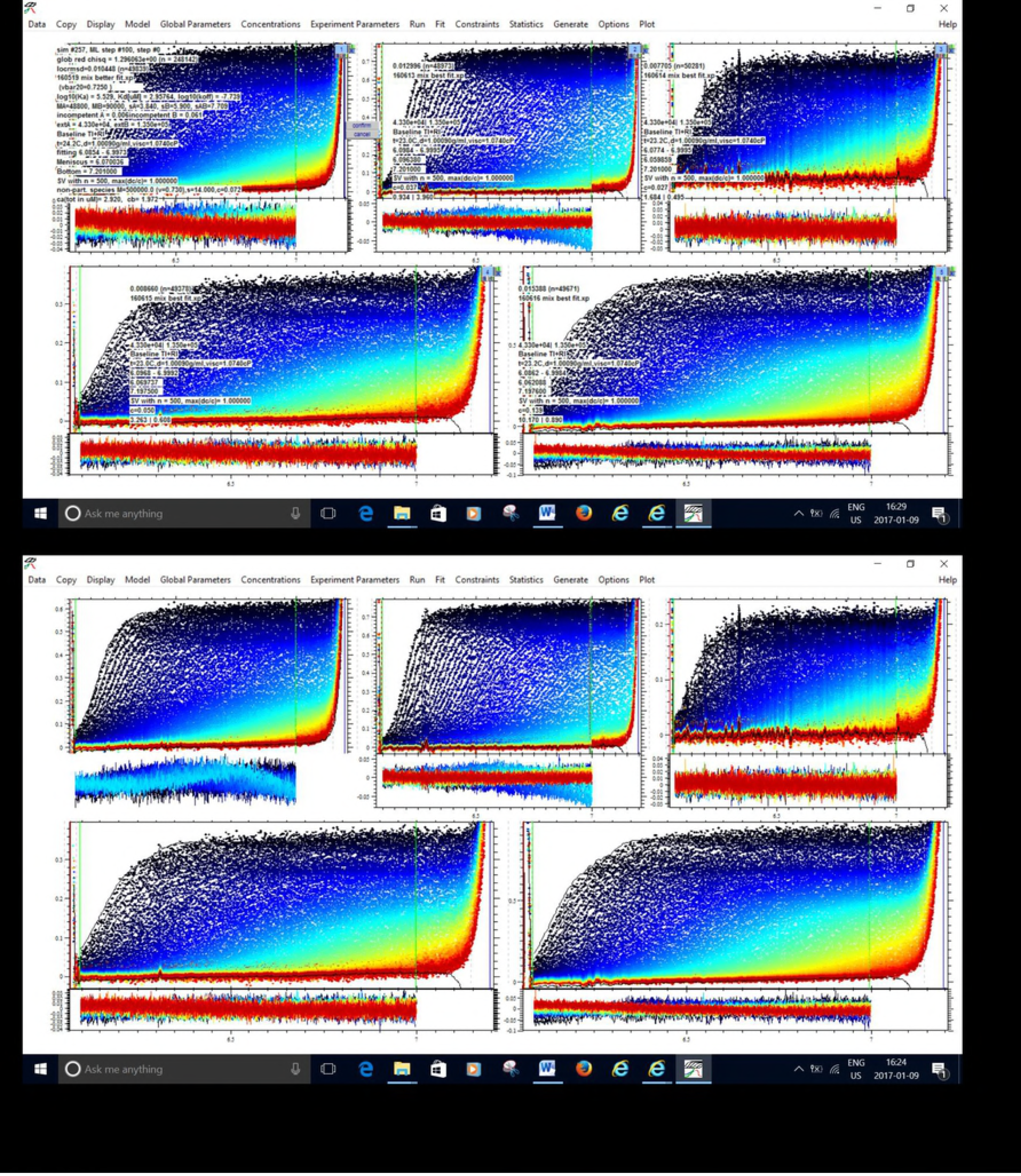
AUC of Str. enolase 137/363 -4, DPgn and the mix of the two proteins. The Sedfit analysis. Lower Panel: The black trace is that of DPgn run separately; the red trace is Str. enolase 137/363 -4. It contains very little octamer. Upper panel: The red trace is the algebraic sum of the two traces in the lower panel. The black trace is that of the actual fit to the data of the mixture of the two proteins. Note the shift to higher s value in the DPgn curve. The two proteins clearly bind to one another when the major Str. enolase species is monomeric.

The criteria for deciding whether binding occurred were based on both the integrated s value of the DPgn peak and the actual position of the peak. If the peak and the integrated position increased in value in the presence of Str. enolase compared to its absence, we assumed it was because binding of something had occurred to cause the shift. If the s value of the DPgn in the mixture did not change, we concluded that no binding had occurred. All the samples behaved similarly to the data of Figs 7 and S6 and all showed binding except Str. enolase DB no tag, Str. enolase DB and other forms that contained only octamers. The data for all the other Str. enolase forms binding to DPgn are not included in this presentation. They are available on request but they agree with the results obtained by DPI.

If one focuses on the positions of the peaks in Figs 6 and S5, it is clear that they do not shift. For Str. enolase DB we can conclude that the protein does not bind to DPgn in solution phase with a Ka of greater than 10^4^ M^−1^ to 10^5^ M^−1^. This contrasts with our DPI results where we clearly saw binding. The Str. enolase DB is octameric. It contains a small amount of large aggregates but no **detectable** monomer (Table 1). We speculate that it is either the aggregates that are selected and bind in the DPI experiment or a vanishingly small amount of combined monomer and oligomers. In either case they are present in such low amounts that we do not see them in the AUC experiments. This allows us to conclude one of the most important aspects of this study: *The fully functional, fully octameric Str. enolase is not the structure that binds to DPgn.*

Str. enolase 137/363 -4 interacting with DPgn (Fig 7) was chosen for illustration because (1) the protein has the largest proportion of monomer of all the truncations and (2) the analyzed data are the most clear cut. The data of the top panel of Fig 7 show that the mixture of the two proteins results in a significant shift to a larger s value of what would have been the plasminogen peak compared to the sum of the fits of the individual species. This view is reinforced when the data are evaluated by DCDT+ (Fig S6).

Fig 7 shows that once again, the shift in the s value for the mixture of the two proteins is significantly greater than the sum of their fits. This data clearly shows that even in solution, we can demonstrate binding of some forms of Str. enolase to DPgn. All of the DPI, LSPR and AUC data are coherent; they indicate that, with the exception of Str. enolase DB no tag and Str. enolase DB, some component in the preparation of Str. enolase 137/363 can, under select conditions, bind DPgn.

The remaining experiments all deal with Str. enolase 137/363 -4 and its interaction with DPgn. This enolase species was chosen for the reasons discussed concerning Fig 7 and S6. We have carried out a series of sedimentation velocity AUC runs in which we have varied the concentrations and ratios of Str. enolase 137/363 -4 to DPgn in the AUC and compared the s value of the resultant peaks of the mix relative to that of the sum of the individual components. For the data of Fig 7, this would represent the peaks occurring at s = 5.21 (6.30 corrected s_20,w_) vs that at 4.70 (5.7 corrected). We reasoned that if the amount of Str. enolase 137/363 -4 bound to the DPgn determined the s value of the complex then we should be able to fit a binding isotherm to the combined data of 137/363 -4 binding to DPgn at different concentrations. (The approach is detailed in the methods section: implementation of Sedphat [39].). Five concentrations of 137/363 -4 and DPgn were chosen and a global fit to 
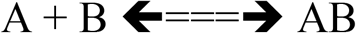
was performed in Sedphat. The results are shown in Fig 8 and are relatively good (locRMSD = 0.01045). The s value returned for the new peak by Sedphat was 7.7 (because we cannot know vbar this is not an s_20,w_). The binding of the Str. enolase 137/363 -4 monomer to DPgn is quite weak. Kd = 3 μM (K_A_ = 3.4E5 M^−1^). By comparison the K_D_s returned by LSPR were 1.3 μM and 2.6 μM. The returned off rate for the AUC global fit, k_off_, was 2E-9. This indicates a very slow off rate which agrees with the data obtained by DPI.

**Fig 8.**
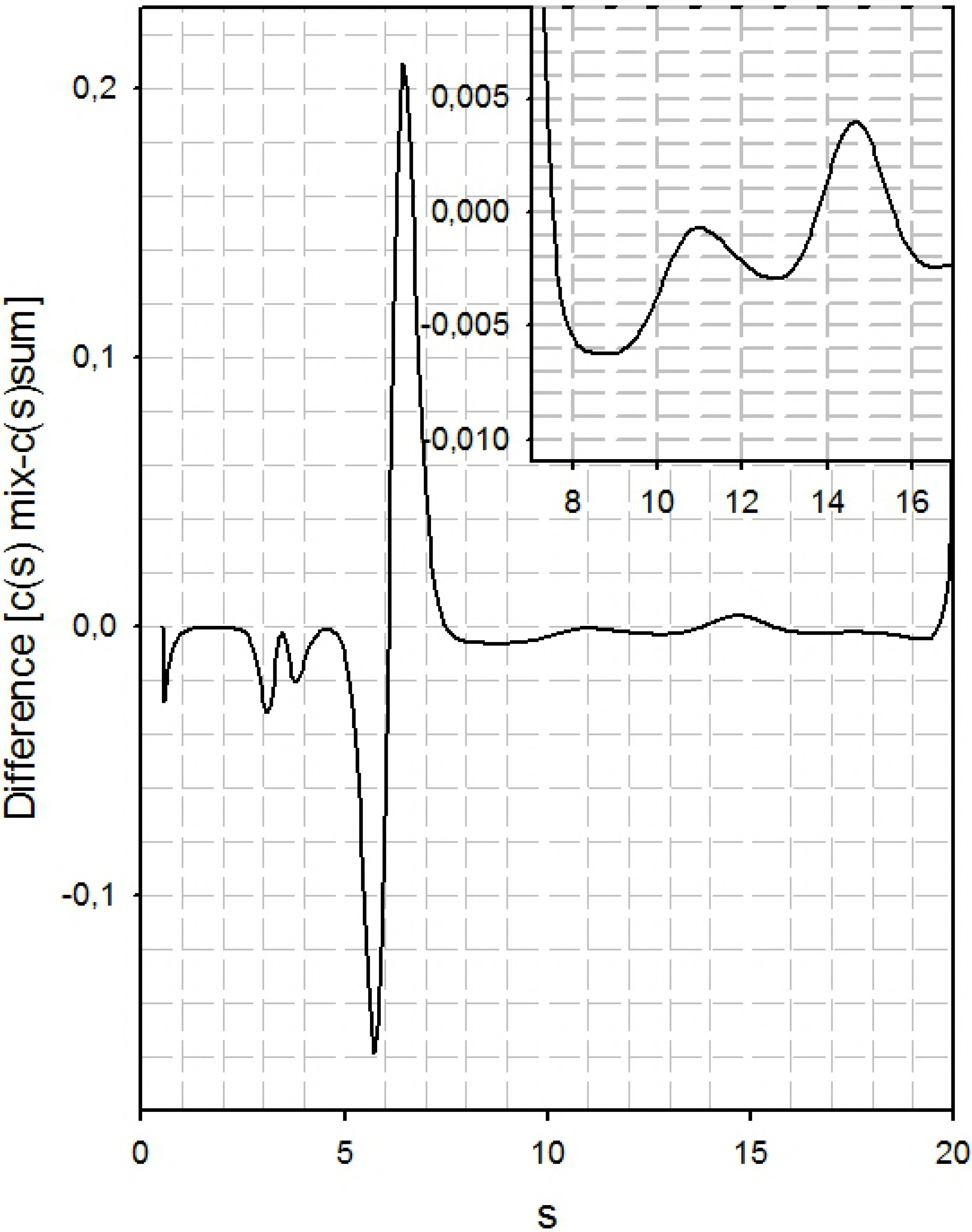
The global fit of 137/363 -4 binding to DPgn: the dissociation coefficient. The two proteins were incubated together and centrifuged. The two sets, upper and lower, show the same data. The lower set shows the data in a way such that the fit lines are visible when zoomed. The upper set shows the fit data and the goodness of fit. The fit data can be easily read when zoomed. The overall locRMSD for the fit is 0.01045. Each panel represents a separate experiment run on a separate day. Str. enolase 137/363 -4 concentrations varied between 1 μM and 11.7 μM; DPgn concentrations varied between 0.43 μM and 4.2 μM.

We then asked the question: can we identify which species in the mix appear and which disappear and does the appearance/disappearance correspond to any known species of Str. enolase 137/363 -4. We approached answering the question by carrying out a subtraction of the sum of the c(s) analysis (137/363 -4 + DPgn) from the actual mixture c(s) analysis. The difference “spectrum” is shown in Fig 9.

**Fig 9.**
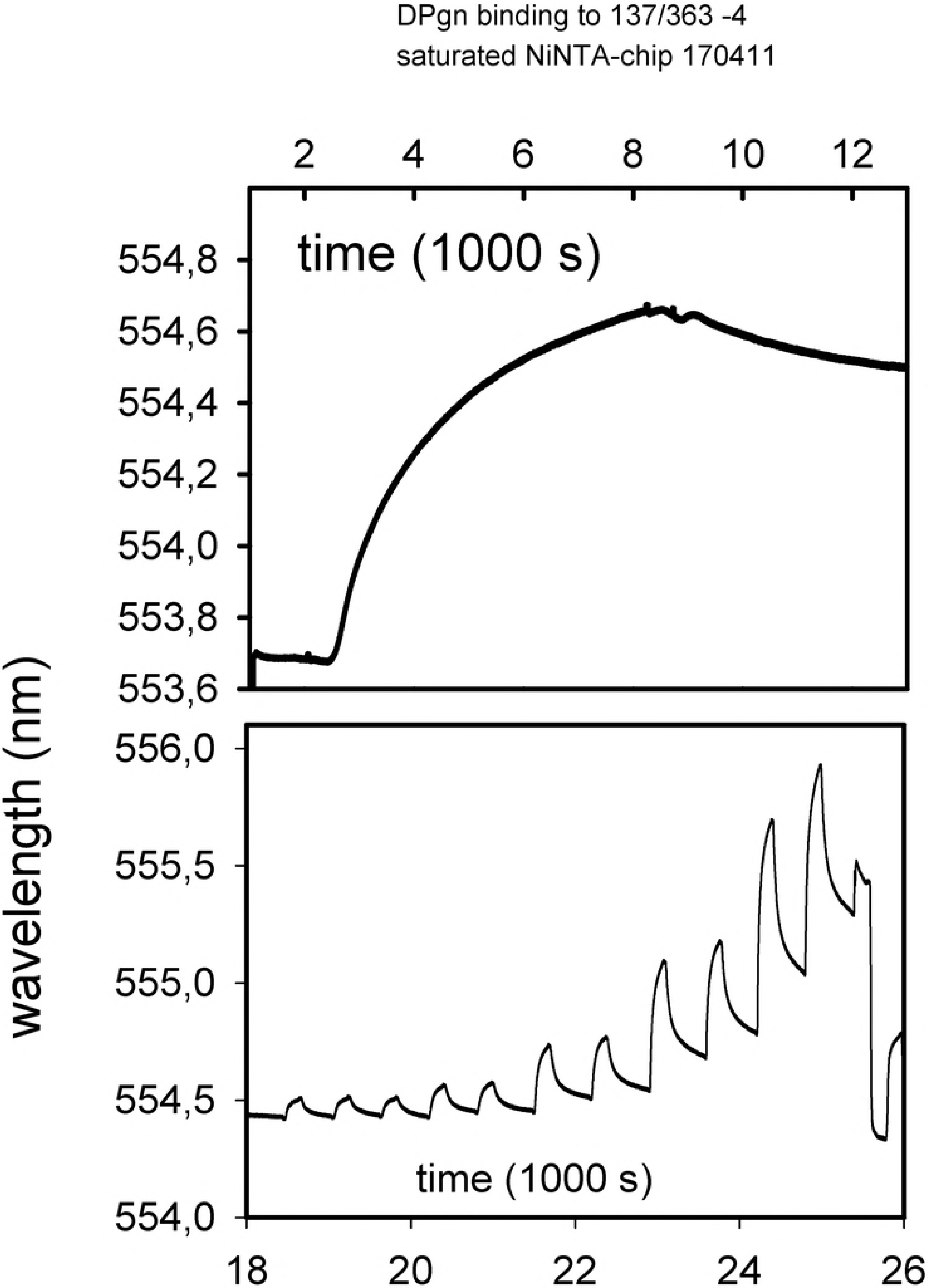
The difference plot emphasizes the loss of components. A plot of the mixture of [Str. enolase 137/363 -4 and DPgn] − [the algebraic sum of the individual c(s) analyses]. The 137/363 -4 concentration was 4.75 μM and DPgn was 1.82 μM, concentrations relatively far from the Ka. The major gain of components from the mix is in the region of s 6.5. The major troughs are at s = 5.7 (DPgn and Str. enolase 137/363 -4?) and at s = 3-4 (Str. enolase 137/363 -4). The eye would indicate that the loss of Str. enolase 137/363 -4 is substantially less than the loss of dimers + DPgn. It must be noted that the 280 nm extinction coefficient of the Str. enolase monomer is only 1/3 that of DPgn. The inset emphasizes the difference spectrum between s values 8 and 16.

The following features about the difference spectrum are notable. The trough at s = 3-4 is probably the result of the complex formation by the monomer of 137/363 -4. The trough at s = 5.7 represents the DPgn and possibly the Str. enolase 137/363 -4 dimer. The complex has moved to s = 6.5. The broad trough centered at s ~ 8 may represent two species neither of which correspond to Str. enolase 137/363 -4 monomers, dimers (s = 6) or tetramers but could be a mix of rapidly exchanging species. One of the characteristics of DPgn is that it undergoes a closed to open conformational transition when DPgn binds both large and small ligands, such as 6-aminohexanoic acid or Str. enolase. The conformational change is large. The s_20,w_ value of the closed conformer is about 5.9 at the temperature used in this study whereas the open conformation has an s_20,w_ of about 4.8 [34]. Clearly, if the DPgn binds a ligand to the open form the s value of the ensuing complex can be considerably smaller than one might anticipate on the basis of assuming a spherical complex.

For the AUC experiments we emphasize that we interpret the experimental result as indicating weak binding but a very slow off rate. This interpretation is similar to that derived from DPI (where we do not have K_A_s) and LSPR (where we do). The association coefficients calculated from LSPR and AUC are quite similar. In this case, surface and solution chemistry yield conclusions which are the same.

In conclusion, we have shown, by DPI, that neither the two carboxy-terminal lysines nor the internal lysines at positions 252 and 255 are required for binding. With the exception of the enolase lacking an N-terminal his-tag, all the enolases examined bound to the DPI chip and then bound DPgn. This was true whether the initial enolase was ≥80 % octameric, or ≤5 % octameric or mainly large aggregates. With the DPI, we do not know what forms of enolase bind to the chip. We also do not know which species on the chip are binding the DPgn. We emphasize that when considering a protein binding non-specifically to a chip, it is difficult to know just which species or subspecies is binding. *Can we be reasonably certain of which species is binding in the AUC experiments? We can be reasonably certain of which species do not bind*. Catalytically active octamers do not bind. That leaves monomers, dimers, trimers and tetramers. Considering the very low percentage of dimer in the Str. enolase 137/363 -4 preparations (Fig 9 bottom panel) it would appear as though the major species for the truncations binding to DPgn is the monomer. It would be interesting to work with enolase on the surface of the gram positive *Streptococcus pyogenes*, perhaps to cross link surface proteins in order to determine their oligomeric status. Unfortunately, this laboratory has neither the technical skills nor the background to work with pathogens.

